# Temple PPG Morphology Demonstrates a Stronger Cardiovascular Age Signal Than Wrist Sites

**DOI:** 10.64898/2026.08.19.745616

**Authors:** Deland Liu, Anirban Dutta, Sachin Nadig

## Abstract

The features of the PPG (photoplethysmography) morphology are known to reflect age-related cardiac and vascular changes. In most contemporary wearables, PPG signals are acquired from distal sites such as the wrist and finger. The superficial temporal artery (STA), accessible at the temple region, is reached via a shorter arterial path from the aortic root than the radial circulation, and may therefore carry hemodynamic and aging information with less distance-dependent attenuation. We hypothesized that the morphology of the PPG at temple region (STA) would show stronger and more numerous age correlates than the PPG at the wrist. To test this, we extracted a common set of 89 pulse-morphology features, spanning raw-waveform timing/amplitude/area measures, ratios among them, derivative-based ratios, and spectral harmonic-ratio features. We compared an in-house temple-worn device which has PPG as one of the sensors, with a publicly available Microsoft Aurora-BP wrist-worn PPG dataset, and tested each feature’s association with age. We identified 14 robust age correlates at the temple region, compared to 3 at the wrist. The temple’s correlates spanned multiple morphological categories and showed a larger age-association than at the wrist. These results support the hypothesis that the temple region may be a more robust PPG measurement site than the wrist to extract age-related cardiovascular information, which motivates further investigation of temple-based cardiovascular sensing.

## 1 Introduction

The aging of the cardiovascular system is accompanied by progressive stiffening of the large elastic arteries, altered propagation and reflection of pressure-flow waves, endothelial dysfunction, and remodeling of the microvascular circulation [38]. These changes modify both the transmission of pulsatile energy through the arterial tree and the morphology of the peripheral arterial pulse. Carotid-femoral pulse wave velocity (cfPWV) is the non-invasive reference measure of aortic stiffness and is independently associated with cardiovascular risk [40, 39, 36]. However, cfPWV requires measurements from spatially separated arterial sites and controlled acquisition procedures, which limits its suitability for continuous or longitudinal wearable monitoring. Photoplethysmography (PPG) can acquire the arterial pulse optically using compact wearable sensors and has therefore attracted considerable interest as a scalable approach to vascular phenotyping [7, 9]. The information contained in a PPG pulse extends beyond the heart rate. Its systolic rise, peak morphology, inflection points, dicrotic region, pulse width, area, and first- and second-derivative landmarks are shaped jointly by ventricular ejection, arterial compliance, vascular impedance, pressure-flow wave transmission, wave reflection, and the local vascular bed interrogated by the optical sensor. Consistent with this physiology, indices derived from the digital volume pulse have been associated with stiffness of the large arteries, e.g. the classical digital volume pulse stiffness index correlated with cfPWV at *r* = 0.65 [27]. Age-related changes in systolic rise and attenuation of the dicrotic region have also been demonstrated directly using multi-site PPG [2]. Therefore, the morphology of PPG contains biologically meaningful information on cardiovascular aging. As illustrated in Figure 1, temporal and distal arterial pathways produce distinct PPG morphologies. The STA waveform retains multiple systolic features whose timing and relative amplitude are sensitive to arterial transmission and wave interaction. Because arterial stiffness, pulse wave velocity, and the timing of reflected components change systematically with aging, these features provide a plausible morphological substrate for age-related vascular phenotyping [22, 25, 8].

**Figure 1.**
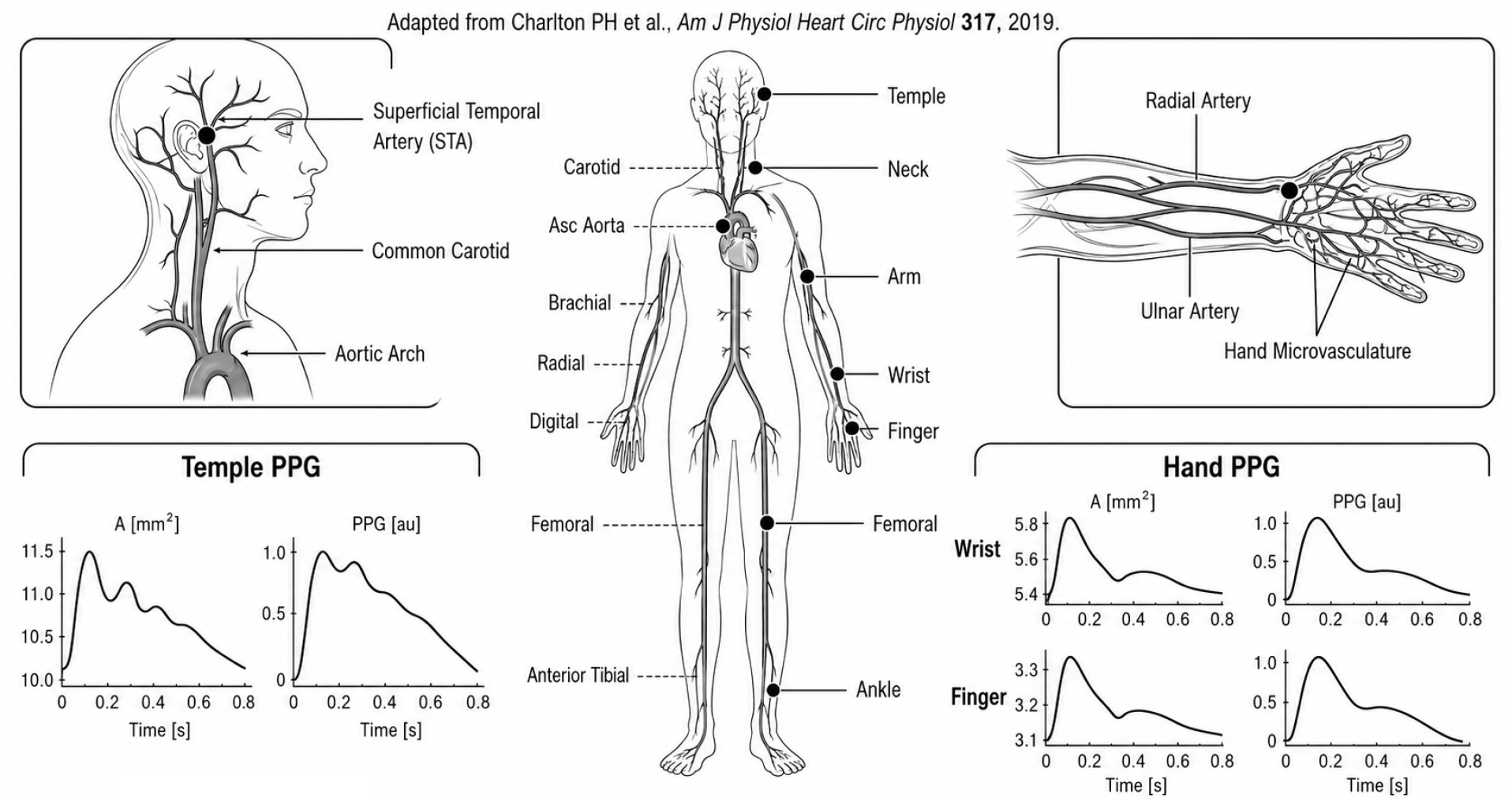
Anatomical and pulse-wave rationale for temple versus distal PPG measurement. The superficial temporal artery (STA) provides a superficial optical target within the carotid arterial territory whereas wrist and finger PPG are measured after propagation through the subclavian–brachial–radial arterial pathway and distal microvasculature. Simulated pulse waves adapted from Charlton *et al*. [8] illustrate pronounced site-dependent differences in waveform morphology. In particular, the STA PPG exhibits a complex multi-phasic systolic contour with secondary peaks, consistent with the superposition of forward and returning wave components and site-dependent arterial transmission whereas the distal wrist and finger waveforms exhibit different peripheral morphologies. These secondary features should be interpreted as reflection and propagation sensitive morphology.

Most contemporary wearable PPG is acquired at distal sites, particularly the wrist and finger, where increasingly sophisticated feature-based machine learning and deep learning models have achieved strong population-level performance. However, substantial inter-individual uncertainty remains because distal PPG morphology is influenced by multiple physiological and site-specific processes and does not uniquely identify the underlying vascular state. In a study of 752 adults aged 20–89 years, direct convolutional neural network analysis of the PPG waveform yielded a mean absolute error (MAE) of 8.1 years (*r* = 0.61), and *R*^2^ = 0.37 for the estimation of chronological-age [33]. Moreover, despite the fact that the Apple Watch wrist PPG achieving excellent chronological-age accuracy (MAE ~2.4 years) [28], our rationale is that the temple/STA PPG may encode aging in richer, cardiovascular, physiologically interpretable waveform features, addressing mechanistic observability rather than outperforming wrist age prediction. We raise a measurement question that has received considerably less attention than model architecture i.e. does the anatomical site at which PPG is acquired constrain the cardiovascular information that remains observable in the pulse waveform?

A PPG waveform is a site-specific optical output of the distributed cardiovascular system. During propagation, pressure and flow waves are modified by arterial geometry, viscoelasticity, branching and wave reflection [42, 8]. The measured waveform is further shaped by the local vascular bed and optical sampling volume [37]. Accordingly, PPG morphology differs significantly between anatomical sites [2, 17]. Wrist PPG reflects central hemodynamics after transformation by the upper limb arterial pathway and local cutaneous circulation [43]. In contrast, temple PPG targeting the superficial temporal artery (STA) samples a superficial external carotid branch arising from the common carotid pathway and may therefore provide a distinct, more proximal richer observation of pulse wave morphology [10, 8]. This distinction is important because increasing algorithmic complexity can exploit nonlinear information retained in a PPG waveform but cannot uniquely recover physiological degrees of freedom that are non-identifiable from the acquired measurement i.e. learned solutions may instead depend on population priors and become unstable when the inverse problem is poorly constrained [34, 3].

These considerations motivated our investigation of the temple, where the STA provides an accessible pulsatile target for optical sensing. The STA is a terminal branch of the external carotid artery and represents a more proximal arterial measurement site than the radial or digital circulation although its waveform remains shaped by local external carotid branching and temple vascular bed properties. Previous studies have demonstrated that reproducible pulsatile hemodynamic signals can be acquired from the STA using PPG-based volume-clamp methods and arterial tonometry [10, 6]. This raises the central question addressed here i.e. *does STA-targeted temple PPG express age-related cardiovascular changes through a broader and more strongly organized set of interpretable pulse morphology features than distal wrist PPG?*

The suitability of this location for optical pulse sensing is supported by previous STA-targeted PPG systems developed for continuous monitoring of arterial pressure [10]. Although the STA volume clamp method used near infrared illumination, green wavelengths preferentially sample superficial pulsatile vasculature [37]. Moreover, at the temple, this includes the richly vascularized cutaneous territory supplied by STA branches, whose microvascular density, endothelial function and vasomotor reserve are themselves altered with aging, potentially adding an age-sensitive cutaneous microvascular component to the more proximal STA pulse wave information [20, 4, 23]. Green PPG has also shown greater robustness to motion related error than infrared PPG in wearable measurements [24]. Importantly, recent multisite measurements demonstrated high-quality reflective PPG at the temple, with green illumination producing the strongest pulsatile component across nearly all investigated anatomical sites [32]. Moreover, the superficial course of the STA provides a favorable optical target where direct placement of a reflectance sensor on the STA produced significantly larger plethysmographic pulsations than forehead placement [31]. The physiological rationale for STA-territory PPG extends beyond technical accessibility. Since the external and internal carotid arteries originate from a shared common carotid input, a pulse measured in the STA samples a branch of the carotid system before the additional transformations imposed by the upper limb arterial tree. The external and internal carotid circulations are anatomically and functionally distinct after bifurcation, nevertheless, carotid territory pulse transmission is of particular physiological interest because aging of the proximal aorta alters the impedance relationship between the aorta and carotid circulation, thus modifying the transmission of pulsatile energy toward the head [29, 14].

We hypothesize that vascular aging may be expressed not only through changes in individual PPG indices but also through the multidimensional structure of age-sensitive pulse wave morphology. Conventional analysis frequently reduce the waveform to one stiffness, reflection, augmentation, or second-derivative aging index. Such a scalarization is convenient but physiologically restrictive. A pulse contains several partially coupled domains where timing descriptors characterize the pulse wave organization of systole and diastole, amplitude and area descriptors characterize the relative expression of waveform components, derivative features emphasize changes in contour and inflection structure, and normalized ratios reduce some forms of inter-subject amplitude variability [8]. These variables can be grouped into mechanistically motivated feature families according to whether they are expected to be predominantly influenced by cardiac ejection, vascular transmission, and reflection or their interaction. We consider these categories as physiological interpretations because no PPG feature on the surface is uniquely generated by a single cardiovascular subsystem.

Large artery stiffening, changes in ventricular-arterial coupling, altered peripheral impedance, and changes in wave reflection timing do not necessarily progress in parallel. An informative vascular waveform may therefore contain several age-sensitive but non-redundant morphological axes. In contrast, if many nominally different PPG features are highly collinear, apparent multiplication of biomarkers does not represent additional physiological information. The covariance structure of the feature space is consequently itself informative where the focus of variance into a small number of components, pairwise feature dependence, and the stability with which individual descriptors are associated with age provide complementary measures of how much independent morphological information is recoverable at a measurement site.

Therefore, the present study tests this site of measurement hypothesis for PPG-based cardiovascular phenotyping. We characterize the fiducials, temporal, amplitude, area, and derivative features of the pulse waveform acquired from the temple/STA territory and compare their age associations and covariance structure with those obtained from wrist PPG. We evaluate whether age sensitivity is distributed across multiple families of physiological characteristics and whether those associations represent independent or redundant manifestations of the same waveform change. We hypothesize that the measurement of the more proximal carotid territory will retain a broader set of age-sensitive morphological characteristics and that its feature space will exhibit greater effective dimensionality than the PPG of the distal wrist. Importantly, chronological age is used here as an initial biological ordering variable rather than as a direct measure of arterial stiffness. The resulting phenotype is therefore interpreted as an age-sensitive vascular waveform feature providing a mechanistic basis for subsequent validation against independent measures of vascular aging.

## 2 Results

### 2.1 Short-separation NIRS-derived PPG shows cardiac and vascular features correlated with age at the approximate temple region

As shown in Table 1 and Table 2, the short-separation NIRS-derived PPG – a channel normally discarded as a noise regressor to remove systemic and superficial signal from long-separation channels [15] – collected in the approximate temple region carries pulsatile cardiac and vascular features that correlate with age. At 830 nm, `AUCsys`, a cardiac feature based on the categorization of the synthetic pulse waveforms in Table 6, remained significant after correction for multiple comparisons (|*r*| = *−*0.852, *p*_corrected_ = 0.0195); no feature survived correction for multiple comparisons at 690 nm.

**Table 1:**
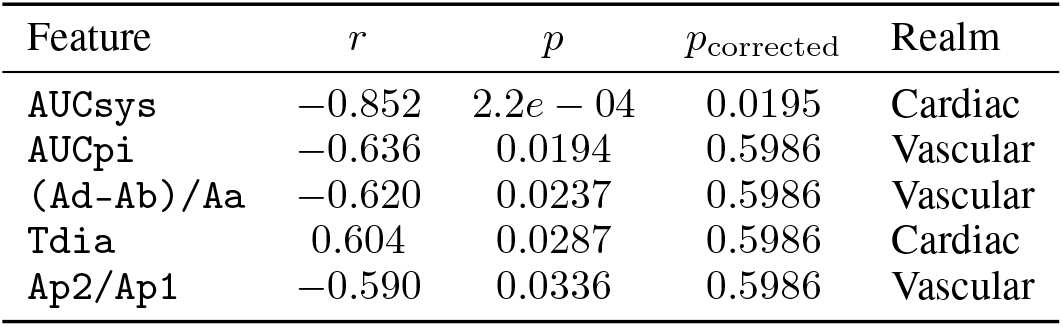
Age correlates significant before multiple-comparisons correction (*p <* 0.05) at the approximate temple region (N=13). *r* is the Pearson correlation coefficient between the feature and age; *p* is the uncorrected (raw) *p*-value from that correlation; *p*_corrected_ is the Benjamini–Hochberg false-discovery-rate-corrected *p*-value across all features tested. Realm indicates whether the feature’s variation is cardiac-, vascular-, or cardiovascular-dominant, per the categorization on synthetic-database. Wavelength: 830nm.

| Feature | $r$ | $p$ | $p_{\text{corrected}}$ | Realm |
| --- | --- | --- | --- | --- |
| AUCsys | -0.852 | $2.2e - 04$ | 0.0195 | Cardiac |
| AUCpi | -0.636 | 0.0194 | 0.5986 | Vascular |
| (Ad-Ab)/Aa | -0.620 | 0.0237 | 0.5986 | Vascular |
| Tdia | 0.604 | 0.0287 | 0.5986 | Cardiac |
| Ap2/Ap1 | -0.590 | 0.0336 | 0.5986 | Vascular |

**Table 2:** Age correlates significant before multiple-comparisons correction (*p <* 0.05) at the approximate temple region (N=13). *r* is the Pearson correlation coefficient between the feature and age; *p* is the uncorrected (raw) *p*-value from that correlation; *p*_corrected_ is the Benjamini–Hochberg false-discovery-rate-corrected *p*-value across all features tested. Realm indicates whether the feature’s variation is cardiac-, vascular-, or cardiovascular-dominant, per the categorization on synthetic-database. Wavelength: 690nm.

| Feature | $r$ | $p$ | $p_{\text{corrected}}$ | Realm |
| --- | --- | --- | --- | --- |
| AUCpi | -0.725 | 0.0051 | 0.4521 | Vascular |
| (Ad-Ab)/Aa | -0.682 | 0.0103 | 0.4563 | Vascular |
| AUCsys | -0.614 | 0.0254 | 0.6168 | Cardiac |
| Tdia | 0.567 | 0.0431 | 0.6168 | Cardiac |
| Tdw50 | 0.557 | 0.0479 | 0.6168 | Cardiac |

Across both wavelengths, the significant features before correction ranged from moderate to very strong correlations with age (*r* ranging from 0.557 to 0.852) and covered both the cardiac and vascular realms in roughly equal measure, without a feature falling into the mixed cardio-vascular category (Table 6). These features spanned three distinct classes of pulse morphology: raw pulse-area measures (`AUCsys`, `AUCpi`), diastolic timing (`Tdia`, `Tdw50`), and second- and third-derivative amplitude ratios (`(Ad-Ab)/Aa`, `Ap2/Ap1`). With the exception of `AUCsys` at 830 nm, none of these associations survived correction for multiple comparisons. Four features (`AUCpi`, `(Ad-Ab)/Aa`, `AUCsys`, and `Tdia`) were significant before correction at both wavelengths with consistent sign.

### 2.2 Dedicated PPG at the temple shows multiple robust age correlates across cardiac and vascular features

Of 89 candidate pulse-morphology features extracted from resting-state PPG recorded directly at the temple in an adult cohort, 14 were identified as robust age correlates (Figure 2a). The median |*r*| for these 14 features ranged from a moderate correlation (*r* = 0.229) to a strong correlation (*r* = 0.679). Of the 14, 4 features – covering the cardiac, vascular and cardio-vascular realms – were positively correlated with age, and 10 features – covering the cardiac and vascular realms – were negatively correlated. The Benjamini–Hochberg-corrected selection probability in 1000 bootstrap iterations ranged from 0.3% to 70.0% (Table 3). The median regression slopes of the features against age ranged in magnitude from 0.0012 to 0.0592 units per year.

**Table 3:**
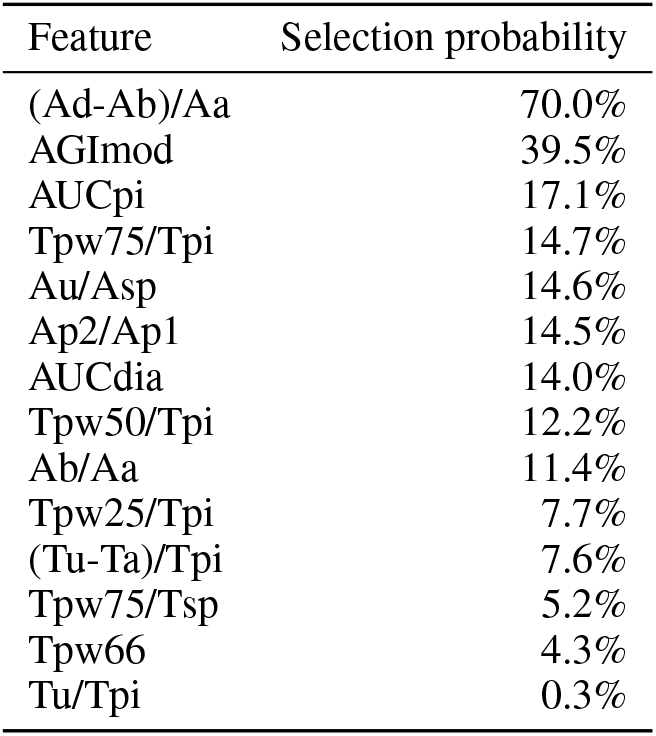
Bootstrap-FDR selection probability of robust PPG-derived age correlates — temple region. Selection probability is the fraction of 1000 age-band-stratified bootstrap iterations in which a feature’s correlation with age was significant after Benjamini–Hochberg FDR correction (*α* = 0.05).

| Feature | Selection probability |
| --- | --- |
| (Ad-Ab)/Aa | 70.0% |
| AGImod | 39.5% |
| AUCpi | 17.1% |
| Tpw75/Tpi | 14.7% |
| Au/Asp | 14.6% |
| Ap2/Ap1 | 14.5% |
| AUCdia | 14.0% |
| Tpw50/Tpi | 12.2% |
| Ab/Aa | 11.4% |
| Tpw25/Tpi | 7.7% |
| (Tu-Ta)/Tpi | 7.6% |
| Tpw75/Tsp | 5.2% |
| Tpw66 | 4.3% |
| Tu/Tpi | 0.3% |

**Figure 2.**
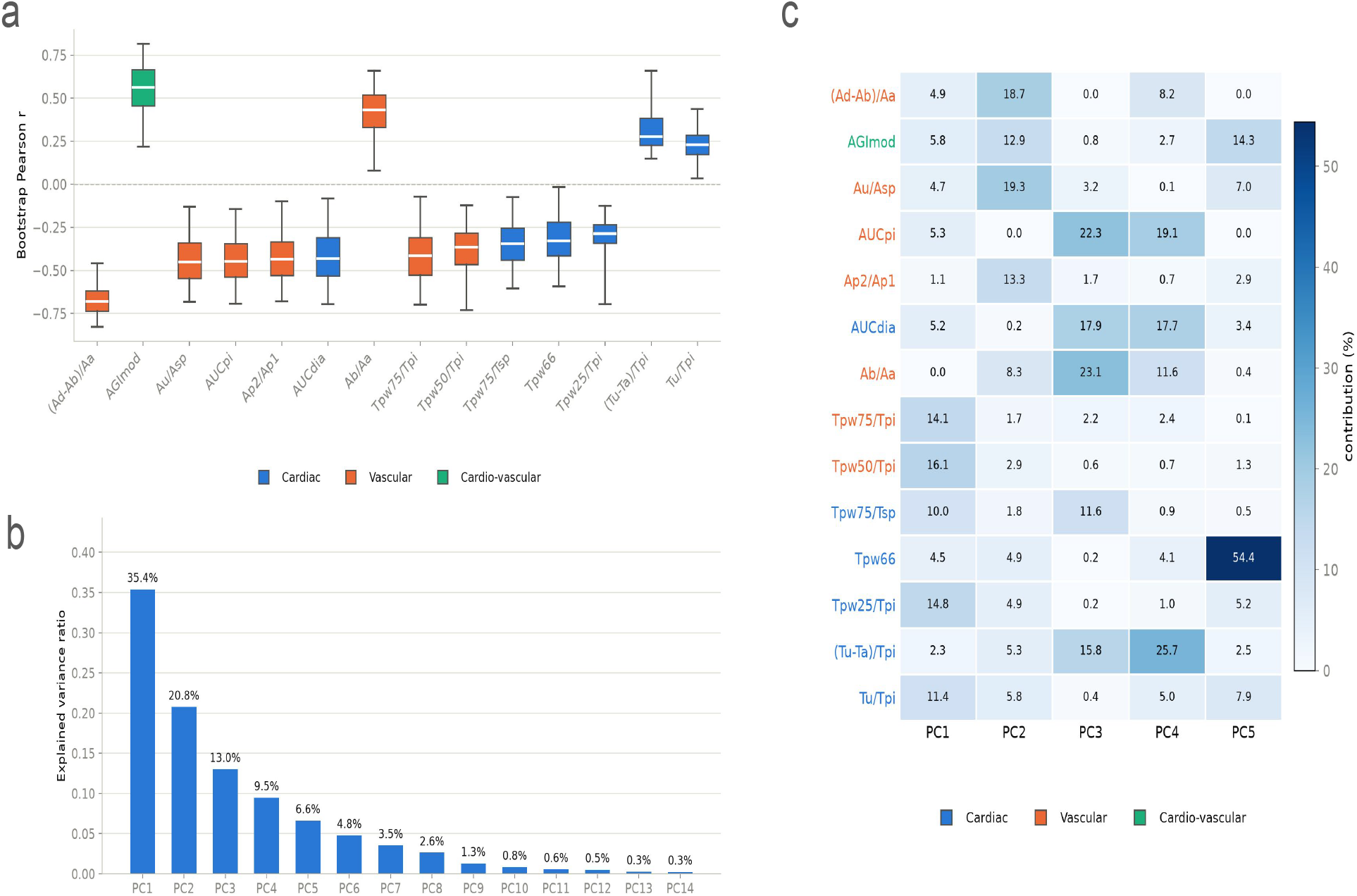
Robust PPG-derived age correlates and their principal-component structure at the temple site. **(a)** Bootstrap distribution (1000 iterations, balanced stratified bootstrap by age band) of the Pearson correlation (r) with age for each PPG-derived feature classified as a robust age correlate at the temple site (N = 122). The box spans the 25th–75^th^ percentile of its bootstrap r distribution, the white line marks the bootstrap median, and the whiskers mark the 95% CI. Box color indicates the feature’s physiological realm classification (Cardiac, Vascular, or Cardio-vascular). **(b)** Scree plot of the variance explained by each principal component of a PCA fit on the robust features across the full complete-data cohort (N = 122). **(c)** Heatmap of each robust feature’s percent contribution to the leading principal components, restricted to the PC1–PC5 needed to reach 80% cumulative explained variance; row labels are colored by the same realm classification as in **(a)**.

The vascular-realm ratio feature `(Ad-Ab)/Aa` was the strongest and most consistently identified correlate, combining the highest correlation magnitude (*r* = *−*0.679) with the highest selection probability (70.0%) of any feature tested. The cardio-vascular ratio feature `AGImod` was the strongest positive correlate (*r* = 0.561, selection probability 39.5%).

The principal component analysis of the 14 robust correlates yielded explained-variance ratios of 35.4%, 20.8%, 13.0%, 9.5% and 6.6% for the first five components (cumulative 85.3%; Figure 2b). For each of these components, the highest load features spanned the cardiac and vascular realms (as categorized in Table 6) and included both amplitude-ratio and pulse-timing-ratio feature types, rather than grouping by realm or feature type (Figure 2c). No single feature dominated the loading of a component, with one exception: `Tpw66` accounted for 54.4% of PC5, a component that explained 6.6% of the total variance.

### 2.3 PPG at the wrist shows fewer and weaker age correlation

Applying the same 89-feature panel to resting-state PPG recorded at the wrist identified 3 robust age correlates, compared to 14 at the temple region (Figure 3a). The median |*r*| across these 3 features ranged from 0.383 to 0.413 (moderate), and the selection probability ranged from 3.7% to 10.1% (Table 4). The median regression slopes ranged in magnitude from 0.0010 to 0.0080 units/year. All 3 correlates were negatively signed and spanned three separate realms: cardiac (`Ad/Aa`), cardio-vascular (`Au/Asp`), and vascular (`Tpw50/Tpi`), as categorized in Table 7.

**Table 4:** Bootstrap-FDR selection probability of robust PPG-derived age correlates — wrist region. Selection probability is the fraction of 1000 age-band-stratified bootstrap iterations in which a feature’s correlation with age was significant after Benjamini–Hochberg FDR correction (*α* = 0.05).

| Feature | Selection probability |
| --- | --- |
| Ad/Aa | 10.1% |
| Au/Asp | 7.3% |
| Tpw50/Tpi | 3.7% |

**Figure 3.**
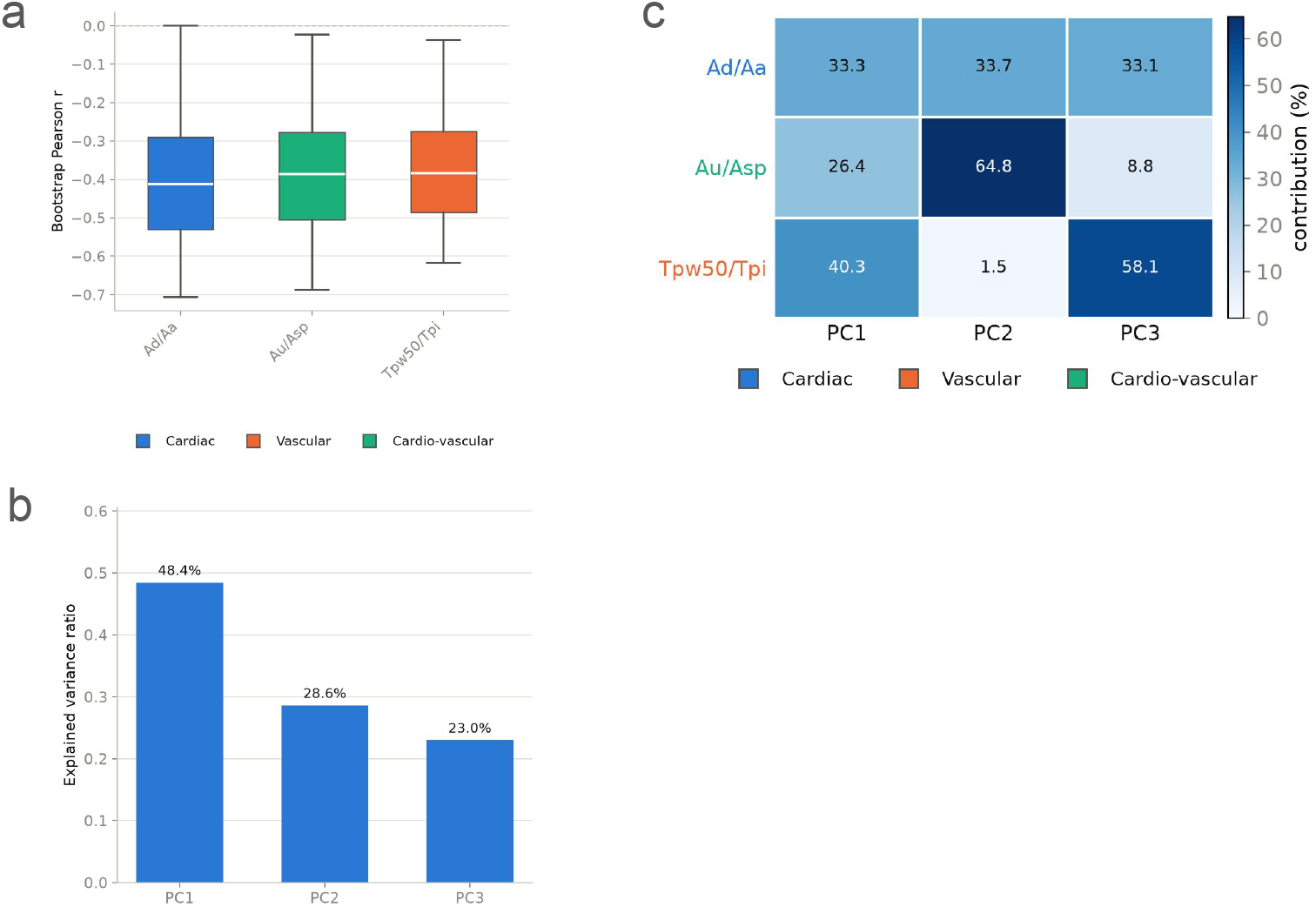
Robust PPG-derived age correlates and their principal-component structure at the wrist site. **(a)** Bootstrap distribution (1000 iterations, balanced stratified bootstrap by age band) of the Pearson correlation (r) with age for each PPG-derived feature classified as a robust age correlate at the wrist site (N = 239). The box spans the 25th–75^th^ percentile of its bootstrap r distribution, the white line marks the bootstrap median, and the whiskers mark the 95% CI. Box color indicates the feature’s physiological realm classification (Cardiac, Vascular, or Cardio-vascular). **(b)** Scree plot of the variance explained by each principal component of a PCA fit on the robust features across the full complete-data cohort (N = 239). **(c)** Heatmap of each robust feature’s percent contribution to the leading principal components, restricted to the PC1–PC3 needed to reach 80% cumulative explained variance (100% covered, as only 3 principal components exist for 3 robust features); row labels are colored by the same realm classification as in **(a)**.

Table 5 reports signed bootstrap *r* with age at both sites for each feature in the union of robust age correlates identified at either site (15 features total). 11 of 15 features showed a larger |*r*| at the temple than at the wrist; of these, 8 have large effect size with |*d*| *>* 1.5 (`(Tu-Ta)/Tpi`, `AUCpi`, `Ab/Aa`, `AUCdia`, `Tpw66`, `(Ad-Ab)/Aa`, `Ap2/Ap1`, `AGImod`), and a further 3 have 0.3 *<* |*d*| *<* 1.5 (`Tpw75/Tpi`, `Tpw75/Tsp`, `Au/Asp`). Amongst the 4 features left, `Tpw25/Tpi`, `Ad/Aa`, and `Tu/Tpi` showed larger |*r*| at the wrist and `Tpw50/Tpi` showed a negligible, direction-ambiguous difference (|*d*| = 0.078).

**Table 5:** Bootstrap correlations (*r*) with age at the temple and wrist sites, shown for the union of all robust age correlates identified at either site (14 at temple, 3 at wrist, 2 shared between sites; 15 total). Values are bootstrap medians with 95% confidence intervals. Cohen’s |*d*| quantifies the effect-size magnitude of the difference between each feature’s temple and wrist bootstrap distributions.

| Feature | Temple $r$ [95% CI] | Wrist $r$ [95% CI] | Cohen’s $\vert d\vert $ |
| --- | --- | --- | --- |
| (Tu-Ta)/Tpi | 0.279 [0.150, 0.660] | −0.139 [−0.508, 0.268] | 2.644 |
| AUCpi | −0.447 [−0.694, −0.144] | 0.016 [−0.364, 0.435] | 2.520 |
| Ab/Aa | 0.432 [0.079, 0.660] | −0.063 [−0.470, 0.377] | 2.507 |
| AUCdia | −0.432 [−0.696, −0.082] | 0.032 [−0.357, 0.443] | 2.448 |
| Tpw66 | −0.329 [−0.594, −0.017] | 0.025 [−0.383, 0.393] | 1.864 |
| (Ad-Ab)/Aa | −0.679 [−0.829, −0.458] | −0.405 [−0.723, 0.015] | 1.834 |
| Ap2/Ap1 | −0.434 [−0.680, −0.100] | −0.111 [−0.500, 0.306] | 1.761 |
| AGImod | 0.561 [0.218, 0.816] | 0.277 [−0.181, 0.622] | 1.568 |
| Tu/Tpi | 0.229 [0.036, 0.438] | 0.356 [−0.016, 0.683] | 0.841 |
| Tpw75/Tpi | −0.415 [−0.699, −0.072] | −0.286 [−0.608, 0.069] | 0.796 |
| Tpw75/Tsp | −0.346 [−0.605, −0.075] | −0.281 [−0.555, 0.143] | 0.567 |
| Ad/Aa | −0.326 [−0.623, 0.051] | −0.413 [−0.707, −0.000] | 0.492 |
| Au/Asp | −0.451 [−0.682, −0.129] | −0.386 [−0.687, −0.023] | 0.335 |
| Tpw25/Tpi | −0.287 [−0.696, −0.125] | −0.361 [−0.681, 0.032] | 0.195 |
| Tpw50/Tpi | −0.365 [−0.730, −0.121] | −0.383 [−0.617, −0.037] | 0.078 |

`Ad/Aa` was the strongest wrist correlate with the highest correlation magnitude (*r* = *−*0.413) and the highest selection probability (10.1%) of the three. Compared against the strongest temple correlate, `(Ad-Ab)/Aa` (*r* = *−*0.679, 70.0% selection probability; Section 2.2), `Ad/Aa` showed approximately 1.6 times lower correlation magnitude and approximately 6.9 times lower selection probability.

PCA of the 3 wrist correlates yielded explained-variance ratios of 48.4%, 28.6%, and 23.0% for PC1–PC3 (Figure 3b). `Ad/Aa` contributed roughly evenly across all three components (33.3%, 33.7%, 33.1%); `Au/Asp` contributed 64.8% of PC2; `Tpw50/Tpi` contributed 58.1% of PC3 (Figure 3c).

Two of the three wrist correlates, `Au/Asp` and `Tpw50/Tpi`, were also identified as robust age correlates at the temple (Figure 2), with the same sign in both cohorts. Note that age-correlates identified at temple region span timing, area, derivative morphology, whilst wrist only covers timing and derivative morphology.

### 2.4 PPG morphology at the temple shows a potential trend toward greater richness than at the wrist

The mean pairwise |*r*| was computed across all the 89 features common to the temple and wrist panels with complete data at both sites, using a balanced age-band-stratified bootstrap. The bootstrap median of this statistic was 0.372 at the temple and 0.385 at the wrist (Figure 4). The bootstrap distribution of the difference (wrist *−* temple) had a median of 0.012. Cohen’s *d* effect size for this difference approached medium size at 0.451, which may potentially reflect a modest trend toward greater PPG richness at the temple than at the wrist.

**Figure 4.**
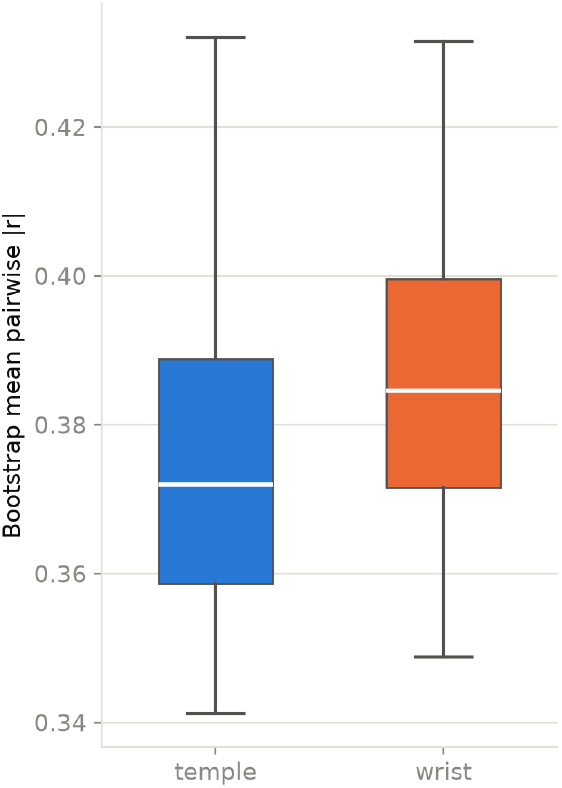
Bootstrap distribution of mean pairwise |*r*| across the 89 features common to both cohorts, temple vs. wrist. Box shows the median and interquartile range (25th–75th percentile) of the bootstrap distribution; whiskers show the 95% CI. Lower mean pairwise |*r*| indicates a less redundant, richer feature set.

## 3 Discussion and Conclusion

The main finding of this study is that the age-related variation in the morphology of PPG was expressed more extensively in the temple region than in the wrist dataset when both were interrogated using the same predefined panel of 89 morphological features. Fourteen features were identified as robust age correlates at the temple, compared to the three at the wrist, and the strongest temple association, `(Ad-Ab)/Aa`, combined a relatively large correlation coefficient (*r* = *−*0.679) with greater bootstrap-FDR selection stability (70.0%) than others feature. The modified AGI, `AGImod`, provided the strongest positive temple association (*r* = 0.561; selection probability 39.5%). Importantly, the temple associations were distributed across derivative morphology, pulse area and timing rather than being restricted to one feature family.

Across the union of age correlates from both sites, the correlation coefficients of most correlates differed between sites with a large effect size, generally favoring a stronger association at the temple. Separately, PPG morphology at the temple showed a slight, suggestive trend toward lower feature redundancy than at the wrist. These observations suggest a measurement site effect on the expression of cardiovascular aging in PPG morphology, although the present between-cohort design does not establish anatomical site as the sole cause of that difference.

The site comparison further suggests that the difference between temple and wrist PPG is not simply an increase in the magnitude of a single age marker but a reorganization of which morphological features express age sensitivity (Table 5). Eight features showed large between-site differences (|*d*| *>* 1.5), spanning timing, pulse-area, amplitude-ratio and second-derivative domains. Particularly notable was `(Ad-Ab)/Aa`, which showed a strong negative association with age at the temple (*r* = *−*0.679, 95% CI [*−*0.829, *−*0.458]) but a weaker and less stable association at the wrist (*r* = *−*0.405, 95% CI [*−*0.723, 0.015]). Several other features, including `AUCpi`, `AUCdia`, `Ab/Aa`, and `Ap2/Ap1`, similarly exhibited larger age effects at the temple whereas some timing features even changed the direction of their age association between sites. Taken together, these findings are consistent with anatomical location acting as part of the physiological observation operator such that cardiovascular aging is expressed differently after propagation through distinct arterial pathways and local vascular beds. The contribution of the local cutaneous microcirculation should therefore be examined explicitly in future studies to disentangle site-specific vasomotor and microvascular effects from aging markers arising predominantly from larger artery transmission and pulse wave morphology. Our interpretation is also consistent with the site dependent determinants observed in the synthetic pulse wave model where the same morphological feature could be predominantly vascular at the temporal site but predominantly cardiac at the radial site. Because the temple and wrist measurements were obtained from different cohorts and devices, however, these between-site effect-size differences should be considered hypothesis generating rather than evidence that the temple is intrinsically superior. Here, confirmation requires paired, within-subject recordings under matched acquisition conditions.

### 3.1 An early-to-late systolic curvature contrast is the dominant age feature at temple site

The particularly strong behavior of `(Ad-Ab)/Aa` is physiologically plausible. Algebraically,

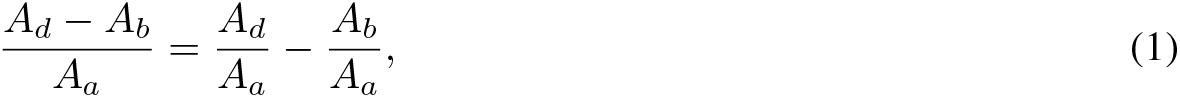

The feature is not the amplitude of a single derivative landmark but a normalized contrast between two temporally separated components of systolic PPG curvature. Under the standardized pyPPG definitions, the *a*-*d* fiducials are typically located during systole with *a* denoting the initial positive second-derivative maximum, *b* the subsequent minimum, and *c* and *d* subsequent extrema before the transition towards the *e* component [16]. These landmarks are interpreted morphologically rather than as unique causal assignments to individual hemodynamic events.

The direction of the association observed here is similar with established aging patterns. In 300 healthy adults aged 19-69 years, the *b/a* ratio progressively increased with age whereas *c/a* and *d/a* decreased [11]. Similar age dependence of second-derivative morphology has been reported in earlier populations [35, 18]. Consequently, the two terms comprising equation 1 move in opposite directions during adult aging with *d/a* decrease while *b/a* increase. Their subtraction can therefore reinforce the age-related transformation of the waveform. This provides a parsimonious explanation for why `(Ad-Ab)/Aa` was more strongly and consistently associated with age than either a single derivative amplitude or the more composite `AGImod` index.

This interpretation is also consistent with the exploratory short separation fNIRS analysis. Although that dataset was small (*N* = 13) and most associations did not survive correction for multiple comparisons, `(Ad-Ab)/Aa` was negatively associated with age at 830 nm (*r* = *−*0.620) and 690 nm (*r* = *−*0.682), with the same direction as in the dedicated temple PPG cohort. Because the short-separation channels were not designed to localize the STA and the sample was underpowered for definitive inference, this result is best regarded as cross-modal and cross-wavelength triangulation rather than independent validation.

### 3.2 The temple phenotype is compatible with wave interaction but is not a direct measure of wave reflection

The early-to-late systolic nature of `(Ad-Ab)/Aa` makes arterial wave transmission a plausible contributor to its age dependence. The arterial pulse observed at any site results from ventricular ejection propagated through a branching, compliant arterial network and modified by distributed impedance discontinuities, local vascular properties, and the superposition of forward and backward traveling waves. Age related increases in arterial wave speed and changes in arterial geometry therefore alter not only pulse timing but also the curvature of the systolic waveform [8, 9].

Charlton *et al*. simulated markedly different PPG morphologies throughout the arterial tree and reported the development of a second systolic peak in carotid flow accompanied by second systolic peaks in STA luminal-area and PPG waveforms [8]. The corresponding limb waveforms underwent different transformations. This result is directly relevant to the present finding because `(Ad-Ab)/Aa` quantifies the relationship between the earlier and later systolic curvature rather than a purely diastolic feature. It provides a mechanistic basis for expecting that the same derivative feature may carry different information when measured after propagation through the carotid-STA pathway than after propagation through the subclavian-brachial-radial pathway.

Our controlled synthetic perturbation analysis strengthens this interpretation. `(Ad-Ab)/Aa` was classified as predominantly vascular at the superficial temporal artery but predominantly cardiac at the radial artery, whereas `AGImod` was classified as cardio-vascular at the temple site and cardiac at the radial site. Thus, the physiological determinants of a nominally identical PPG feature were themselves site-dependent. This is consistent with viewing anatomical location as part of the cardiovascular observation operator i.e. the waveform measured at a surface site represents the interaction between a common upstream excitation and the transfer characteristics of the arterial pathway leading to that site. The result should nevertheless be interpreted within the scope of the model, which perturbed a finite set of cardiac and vascular parameters and does not contain every determinant of in-vivo optical PPG.

Pressure-flow wave-intensity measurements have reported that approximately 27% of incident systolic wave energy was reflected in the radial artery, compared to approximately 17% in the carotid artery [44]. Peripheral pressure augmentation can likewise be substantial. Furthermore, second-derivative landmarks are not direct measurements of forward or backward wave amplitude. Simultaneous carotid pressure and flow measurements have shown that conventional augmentation index and waveform shoulder timing can disagree substantially with wave-reflection indices obtained from wave-intensity and wave-separation analyses [19]. We therefore interpret `(Ad-Ab)/Aa` as *wave-interaction- and propagation-sensitive* rather than solely reflection-specific. Candidate determinants include arterial stiffness and wave speed, arterial diameter and compliance, ventricular ejection, the timing and superposition of returning waves, branching and impedance mismatches, and local vascular-bed properties.

This distinction is especially important in the carotid territory. A backward traveling wave can be measured in the common carotid artery and is modified by changes in downstream cerebral vasomotor tone [5]; however, the STA is an external-carotid branch and should not be treated as a direct measurement of internal carotid or cerebral wave reflection. The external and internal carotid arteries instead share a common upstream carotid input before bifurcation. Age-related changes in aortic stiffness and aorta–carotid impedance matching can modify that shared input and the amount of pulsatile energy entering the carotid circulation [29]. Whether these changes specifically account for the temple PPG phenotype observed here requires simultaneous pressure-flow measurements and cannot be established from PPG morphology alone.

### 3.3 Relation to dicrotic morphology and large-scale wearable aging phenotypes

The present findings can also be placed in the context of recent full-waveform aging studies. In the Apple Heart & Movement Study, Miller *et al*. trained a high-dimensional representation using approximately 20 million 60s Apple Watch PPG recordings and subsequently estimated chronological age from repeated wrist measurements [28]. In healthy held-out participants, the full-waveform model achieved an age-estimation error of approximately 2.4 years, markedly lower than models based only on heart rate and heart-rate variability. Inspection of waveforms associated with progressively older predicted age showed attenuation and eventual disappearance of the dicrotic notch and diastolic peak [28].

This large-scale result is important for interpretation of the present wrist–temple comparison. It demonstrates that distal wrist PPG can contain substantial age-related information when many recordings and a high-dimensional representation are used. Our present findings should therefore raise a different question i.e. whether age-related cardiovascular information is expressed more transparently in physiologically interpretable, low-dimensional morphology at some anatomical sites than at others. In our data, conventional fiducial-based analysis recovered substantially more stable age associations at the temple site whereas large-scale wrist studies demonstrate that age information can also be recovered through learned whole-waveform representations.

The relationship between our derivative findings and dicrotic morphology is potentially informative. The *a*-*d* derivative fiducials occur predominantly during systole, whereas the dicrotic region lies later in the waveform. Thus, `(Ad-Ab)/Aa` may quantify an age-dependent reorganization of systolic curvature that accompanies the broader loss of waveform complexity seen in aging PPGs. Consistent with the clinical relevance of this broader phenotype, Cunningham *et al*. analyzed 169,787 UK Biobank finger PPGs and showed that a continuous measure of dicrotic-notch smoothness was heritable, associated with genetic loci implicated in vascular properties, and stratified incident cardiovascular disease [13]. Together, these studies suggest that vascular aging is expressed across several levels of waveform organization, from second derivative systolic curvature to the macroscopic dicrotic contour and ultimately to high dimensional whole waveform representations.

The dicrotic notch itself should not, however, be equated with a single reflected wave. Its physical origin remains debated, and experimental and modeling work supports contributions from ventricular–arterial dynamics, aortic-valve closure, arterial propagation and wave interaction rather than a unique reflected-wave mechanism [1]. The common physiological theme connecting these observations is therefore *age-related re-modeling of pulse-wave morphology*.

### 3.4 AGImod provides a complementary but more composite aging phenotype

The positive association of `AGImod` with age provides complementary evidence. Modified second-derivative aging indices combine several systolic fiducials and have repeatedly been associated with age and arterial properties. In the Charlton’s computational model, `AGImod` correlated with aortic PWV but sensitivity analysis demonstrated that it was not determined by PWV alone; cardiac and vascular properties also contributed to its variation [8]. This supports its classification as a composite morphology phenotype rather than a direct optical measure of arterial stiffness.

The stronger performance of `(Ad-Ab)/Aa` in the present temple cohort is therefore noteworthy. Whereas `AGImod` combines several second-derivative components, `(Ad-Ab)/Aa` isolates the contrast between two components whose adult age trajectories are known to diverge. Its higher correlation magnitude and substantially greater bootstrap selection probability are consistent with the possibility that this early-to-late systolic contrast provides a cleaner age axis at the temple site. This interpretation remains hypothesis-generating until prospectively tested against independently measured hemodynamic variables.

### 3.5 Multidimensionality of the temple PPG age phenotype

Five principal components were required to account for 85.3% of variance among the 14 robust temple correlates and the principal components received contributions from features classified as cardiac, vascular and cardio-vascular. The age phenotype therefore appears to span several partly coupled waveform dimensions rather than being equivalent to one stiffness or reflection index.

Separate redundancy analysis provides only tentative evidence for greater morphological richness at the temple site. The mean pairwise correlation across the common 89-feature panel was slightly lower at the temple than at the wrist but the bootstrap confidence interval for the between-site difference included zero and the standardized effect was modest (*d* = 0.451). Under the present feature-selection procedure, more age-associated features with larger and more stable effects were recovered from the temple dataset.

### 3.6 Limitations and mechanistic tests

Several limitations are central to interpretation. First, temple and wrist signals were obtained from different cohorts rather than simultaneously from the same individuals. Although correlation analyses used balanced, stratified bootstrapping by age band to ensure same age distribution and sample size in comparisons, several cross-cohort differences remained: sample size, age distribution, skin tone, blood pressure, and sex composition varied between cohorts, as did the native sensors, hardware sampling rates and separation distance. Though both wrist and temple datasets used green LED, their wavelengths differed too: wrist (527 nm - 537 nm), temple (520 nm - 535 nm). In particular, the dedicated temple PPG was acquired at 32 Hz whereas the wrist signal was acquired at 500 Hz before both were processed through a common pipeline. Resampling cannot restore morphological frequency content absent from the lower-rate acquisition, and derivative features are especially sensitive to sampling frequency, filtering and fiducial detection. The persistence of strong derivative associations in the temple data is therefore notable, but it does not remove the possibility of acquisition-dependent bias. In addition, a higher age group (>60 years) was not included in the correlation analyses, in order to facilitate comparison between cohorts whose available age ranges did not otherwise align. This reduces sensitivity to interesting cardiovascular aging insights that might accelerate later in life.

Second, correlation with chronological age is not equivalent to validation against vascular aging. Age also covaries with cardiac function, blood pressure, autonomic regulation, body composition and other determinants of PPG. The terms “vascular” and “cardio-vascular” in this study describe mechanistic classifications obtained from controlled synthetic perturbations and they should not be interpreted as proof of a unique in-vivo physiological origin.

Third, the temple sensor was positioned at the STA territory but the arterial localization was not verified by vascular imaging. The measured optical signal therefore contains pulsatile contributions from the named artery and the surrounding cutaneous vascular bed in proportions that remain unknown. Future studies should use ultrasound or Doppler localization where precise STA attribution is required.

Further, the results with fNIRS in Section 2.1 should be interpreted with care. The signal is only an approximate surrogate for temple PPG, which is derived by averaging two short-separation channels over a broader fronto-temporal scalp region. The cohort is small (N=13) with an uneven age distribution. Therefore, the results should be treated as foundational, exploratory work rather than a basis for direct comparison with the temple and wrist results.

The critical next experiment is consequently a within-subject, same-session comparison using matched optical wavelength, sensor geometry, sampling rate and preprocessing at the temple and wrist. Simultaneous cfPWV and carotid or central hemodynamic measurements would determine whether `(Ad-Ab)/Aa` and `AGImod` track vascular function independently of chronological age. A particularly informative mechanistic experiment would combine PPG with pressure–flow or diameter–flow measurements sufficient for wave-intensity or wave-separation analysis. This would permit direct testing of whether the temple PPG derivative phenotype covaries with backward-wave amplitude or timing, or instead with arterial stiffness, ventricular ejection or other components of the local transfer function.

The existing one-dimensional model provides an immediate complementary route to this test. Independent perturbation of PWV, large-artery diameter, mean arterial pressure, stroke volume and LVET at the radial and superficial temple sites can quantify the site-specific sensitivity of `(Ad-Ab)/Aa`. Demonstrating preferential sensitivity of the temple feature to vascular network perturbations while controlling cardiac parameters would provide a considerably stronger mechanistic bridge between the present human association and arterial wave transmission physiology.

## 4 Conclusion

In the cohorts examined, PPG acquired from the temple/STA territory expressed a broader, larger and more reproducible set of age-associated conventional morphological features than wrist PPG. Nonetheless, these results should be interpreted with caution, as the comparison is done between-cohort rather than with a within-subject design. The dominant temple feature, `(Ad-Ab)/Aa`, is a normalized contrast between early and late systolic second-derivative morphology whose negative age dependence is consistent with the opposing adult age trajectories of *b/a* and *d/a* reported previously. Its site-dependent classification in controlled hemodynamic simulations further suggests that anatomical location modifies the physiological determinants represented by a nominally identical PPG feature.

These findings support a site dependent observability hypothesis where arterial propagation, ventricular-vascular interaction, branching, and wave superposition determine how cardiovascular aging is expressed in local PPG morphology. Large-scale wrist studies show that substantial age information can also be recovered distally using high-dimensional whole-waveform models, and the potential advantage of the temple site may therefore lie in making components of that physiology more directly accessible to interpretable richer morphology-based analysis. Prospective validation within the subject against arterial stiffness and direct wave measurements is required before the temple phenotype can be interpreted as a biomarker of vascular aging.

## 5 Methods

### 5.1 Datasets and demographics

#### 5.1.1 fNIRS dataset at the approximate temple region

The resting-state fNIRS data were obtained from Dataset II of the publicly available multimodal fNIRS dataset of [41]. Data were acquired with a multichannel continuous-wave fNIRS system (CW6, TechEn Inc., MA, USA) that covered the the fronto-parietal region bilaterally. The recording setup included 16 sources, along with 24 long-separation detectors (positioned roughly 3 cm from the source) and 8 short-separation detectors (positioned roughly 1 cm from the source), operating at wavelengths of 690 and 830 nm and sampled at a rate of 50 Hz. The cohort consisted of 14 healthy participants (age 32 *±* 19 years; 7 male/6 female/1 not reported) with no reported neurological or psychological disorders, each seated comfortably and instructed to rest quietly for a nominal 10-minute recording. One subject marked as having low SNR was omitted.

The channel-to-anatomical correspondence was determined using a 10-20 reference file accompanying the probe design. We isolated the short-separation channels nearest Fp2, F6, AF8, and FT8, which cover the frontal and fronto-temple scalp approximately overlying the right temple.

For this analysis, we used the anterior patch of the right hemisphere. Of that patch’s channels, only the short-separation pairs were retained. For each of the two wavelengths, the raw intensity time series from these short-separation pairs were averaged to yield a single per-wavelength waveform per participant. This averaged signal was treated as a surrogate PPG.

#### 5.1.3 Microsoft Aurora PPG dataset at wrist

The wrist-worn PPG data was obtained from the Aurora-BP dataset [30, 26], collected with an optical sensing device (MAX30101) mounted underneath a watch body. Green LED was used. The sampling rate of the data was 500 Hz. We used the subset of participants enrolled in the oscillometric protocol, in which a series of measurements were collected in two controlled in-lab visits at least 24 hours apart. From each in-lab visit, only the seated, stationary measurements were used.

To match the healthy-cohort requirement used for the fNIRS dataset, participants were included only if they reported no diagnosed cardiovascular disease, hypertension, diabetes, arrhythmia, prior heart attack or stroke, heart failure, aortic stenosis, valvular heart disease, or cardiovascular medication use, and self-reported normotensive status. After this exclusion criteria, *N* = 240 participants remained (age range 30-60 years, mean 43.3 *±* 8.9; 125 female, 115 male).

#### 5.1.3 Dedicated PPG dataset at temple region

PPG data were collected from an internal cohort using a wearable PPG optical sensor mounted at the right temple region with medical tape, operating at the green wavelength and sampled at 32 Hz. As with the other two datasets, only healthy participants were included, with no known neurological or psychological disorders. Data were restricted to stationary periods. After this filtering, *N* = 126 participants remained, with ages ranging from 20 to 56 years (mean 29.7 *±* 8.3; 29 female, 91 male, and 6 not reported). The study protocol for this data collection was reviewed and approved by SER-IEC (Approval No. SER-IEC/2026/AP/031, Track No. SER-IEC-NR-2026-025). Written informed consent was obtained from each participant prior to study procedures

#### 5.1.4 Synthetic PPG

Synthetic PPG waveforms were obtained from the Pulse Wave Database (PWDB) of [8], a hemodynamic model simulating arterial pulse waves representative of healthy adults across six age groups (25, 35, 45, 55, 65, and 75 years). At each age, a baseline virtual subject was simulated by setting the model’s cardiovascular input parameters to their age-specific mean values. Around this baseline, 729 subjects per age were generated by independently perturbing the parameters found to most strongly influence pulse-wave morphology at *±* 1 SD from the age-specific mean. This yielded 4,374 virtual subjects in total. Note that the database is not stratified by sex.

Simulated PPG waveforms were extracted at two measurement locations corresponding to the two real PPG datasets described above: the radial artery (wrist) and the superficial temporal artery (temple), sampled at 500 Hz.

### 5.2 PPG preprocessing

PPG data from fNIRs, wrist and temple datasets were bandpass-filtered to 0.5-10 Hz, at their individual sampling rates, using a zero-phase, 4th-order Butterworth filter, to suppress Mayer waves (~0.1 Hz) [21], respiratory waves (~0.2-0.3 Hz) [12] that could be mistaken for a cardiac pulse by the fiducial-point detection algorithm.

For the temple-worn PPG and fNIRS-derived signals, the waveform was sign-inverted prior to fiducial extraction, since both produced the inverse of the conventional PPG orientation in which the systolic peak is an upward deflection. The wrist-worn PPG and synthetic waveforms were already provided in conventional PPG orientation and were not inverted.

Each signal was then segmented into non-overlapping 25s duration windows, and was then resampled to a common target rate of 75 Hz. The signal quality was assessed per window using pyPPG’s own beat-template correlation score [16]. Specifically, a template pulse was constructed from the beats within each window, and each individual beat in a window was scored by its correlation to that template. A global cutoff was computed as the 10th percentile of all beat-level scores pooled across every window and participant in a given dataset. A window was then rejected if more than 50% of its own beats fell below that global cutoff. This procedure rejected 30 of 581 candidate windows (5.2%) for the temple-worn PPG dataset, 64 of 1244 (5.1%) for the wrist-worn PPG dataset, and 0 of 372 (830 nm) and 1 of 372 (690 nm; 0.3%) for the fNIRS-derived dataset.

### 5.3 PPG feature extraction

From each window, fiducial points were identified in the raw pulse waveform and on its first, second, and third derivatives, and pulse-morphology features were computed from these points using the pyPPG toolbox [16], yielding a total of 89 features. Three biomarker families used features computed directly from the raw-waveform fiducial points (pulse onset, systolic peak, dicrotic notch, and diastolic peak) and comprising of timing intervals, amplitude differences, and areas under the pulse curve; ratios computed among these same raw-waveform quantities; and lastly, ratios computed among fiducial points from the first, second, and third derivatives of the waveform. Two additional harmonic-ratio features, the power-spectral-density ratio of the cardiac fundamental frequency to its second and third harmonics, were also computed.

For each window, the power spectral density (PSD) was estimated using Welch’s method (segment length 1024 samples). Spectral peaks were identified on the log-power spectrum using a prominence threshold of 0.2. Among these peaks, the one with the highest power falling within the physiological heart-rate band (0.7-2.0 Hz) was taken as the cardiac fundamental. For the second and third harmonics, a target frequency was computed as twice and three times the fundamental frequency, respectively. The detected peak closest to that target among all peaks was accepted as the harmonic if it fell within a tolerance of 5 frequency bins. Each ratio was then computed as the fundamental peak’s power divided by the matched harmonic peak’s power.

Feature values were computed for each detected pulse within a window and aggregated by taking the median across all pulses in that window, yielding one value per feature per window. Harmonic-ratio features, which are computed once per window directly from its power spectral density rather than per pulse, did not require within-window aggregation. These window-level values were then aggregated a second time by taking the median across all of a subject’s windows, yielding a single value per feature per subject.

### 5.4 PPG feature binning

Each extracted PPG morphology feature was classified as cardiac, vascular, or cardiovascular in Tables 6 and 7 using the synthetic dataset’s experimentally-controlled perturbation design [8].

**Table 6:** PPG feature categorization by cardiac, vascular and cardio-vascular determinant, using synthetic pulse-wave model, at the Superficial Temporal Artery.

| Location | Determinant | Features |
| --- | --- | --- |
| Superficial Temporal Artery | Cardiac | (Tu-Ta)/Tpi, AUCdia, AUCsys, Adn, Adp, Adp/Asp, Aoff, Asp/(Tpi-Tsp), Asp/Aoff, Av/Au, IPA, IPAD, IPR, Ta/Tpi, Td/Tpi, Tdia, Tdw10, Tdw25, Tdw33, Tdw50, Tdw66, Tdw75, Tdw90, Te/Tpi, Tf/Tpi, Tpw10, Tpw25, Tpw25/Tpi, Tpw25/Tsp, Tpw33, Tpw50, Tpw50/Tsp, Tpw66, Tpw75, Tpw75/Tsp, Tpw90, Tsp, Tsp/Asp, Tsp/Tpi, Tsw10, Tsw25, Tsw75, Tsw90, Tsys, Tsys/Tdia, Tu/Tpi, Tw/Tpi, deltaT, second harmonic ratio |
|  | Vascular | (Ac-Ab)/Aa, (Ad-Ab)/Aa, (Tv-Tb)/Tpi, AGI, AGIinf, AI, AUCpi, Ab/Aa, Ac/Aa, Ad/Aa, Ae/Aa, Af/Aa, Ap2/Ap1, Au/Asp, Aw/Au, SC, Tb/Tpi, Tdw10/Tsw10, Tdw25/Tsw25, Tdw33/Tsw33, Tdw50/Tsw50, Tdw66/Tsw66, Tdw75/Tsw75, Tdw90/Tsw90, Tpw50/Tpi, Tpw75/Tpi, Tsw33, Tsw50, third harmonic ratio |
|  | Cardio-vascular | AGImod, Asp, Asp/deltaT, RIp1, RIp2, Tc/Tpi, Tdp, Tpi, Tpp, Tsw66, Tv/Tpi |

**Table 7:** PPG feature categorization by cardiac, vascular and cardio-vascular determinant, using synthetic pulse-wave model, at the Radial Artery.

| Location | Determinant | Features |
| --- | --- | --- |
| Radial Artery | Cardiac | (Ac-Ab)/Aa, (Ad-Ab)/Aa, (Tu-Ta)/Tpi, AGIinf, AGImod, Ac/Aa, Ad/Aa, Adp, Adp/Asp, Ae/Aa, Af/Aa, Asp/(Tpi-Tsp), IPA, IPR, RIp1, RIp2, Tc/Tpi, Td/Tpi, Tdia, Tdp, Tdw10, Tdw10/Tsw10, Tdw25, Tdw66, Tdw66/Tsw66, Tdw75, Tdw75/Tsw75, Tdw90, Tdw90/Tsw90, Te/Tpi, Tf/Tpi, Tpi, Tpp, Tpw10, Tpw25, Tpw75/Tpi, Tpw75/Tsp, Tsys, Tsys/Tdia, second harmonic ratio |
|  | Vascular | (Tv-Tb)/Tpi, AI, AUCdia, AUCpi, Adn, Aoff, Ap2/Ap1, Asp, Asp/Aoff, Asp/deltaT, Av/Au, IPAD, SC, Ta/Tpi, Tdw25/Tsw25, Tdw33/Tsw33, Tdw50/Tsw50, Tpw25/Tpi, Tpw25/Tsp, Tpw50/Tpi, Tpw50/Tsp, Tsw10, Tsw25, Tsw33, Tsw50, Tsw66, Tsw75, Tsw90, Tu/Tpi, Tv/Tpi, Tw/Tpi |
|  | Cardio-vascular | AGI, AUCsys, Ab/Aa, Au/Asp, Aw/Au, Tb/Tpi, Tdw33, Tdw50, Tpw33, Tpw50, Tpw66, Tpw75, Tpw90, Tsp, Tsp/Asp, Tsp/Tpi, deltaT, third harmonic ratio |

For each feature, a separate one-way ANOVA was run for each of the six cardiovascular properties (three for cardiac and three for vascular), testing how much perturbing that property alone (at *−*1, 0, or +1 SD) explained the feature’s variation. Each ANOVA yields a p-value and a partial eta-squared effect size. The three cardiac properties’ partial eta-squared values were summed to give a single cardiac effect-size score per feature, and likewise for the three vascular properties, giving a vascular effect-size score. A feature was labeled Cardiac or Vascular according to whichever summed effect size was larger, unless the ratio of the smaller to the larger summed effect size exceeded 0.8, in which case the feature was labeled Cardiovascular, reflecting that cardiac and vascular properties contribute comparably to that feature’s variance rather than one dominating. This categorization was performed independently for each of the two synthetic measurement locations (radial artery, superficial temporal artery).

### 5.5 Statistical analysis

#### 5.5.1 Age-correlation bootstrap (Temple, Wrist datasets)

For each extracted PPG feature association with age was assessed using a stratified bootstrap rather than a single correlation computed on the full cohort, to reduce sensitivity to the uneven age distribution within each dataset and to put datasets of different sizes on a comparable footing. Subjects were first assigned to three age bands (20-35, 35-45, 45-60 years). In each of 1,000 bootstrap iterations, 8 subjects with replacement from each band (24 subjects total per iteration), and the Pearson correlation coefficient and ordinary least squares (OLS) slope of each feature against age were computed on that sampled set. This yielded, per feature, a bootstrap distribution of 1,000 correlation coefficients and 1,000 slopes, summarized by their median and 95% interval (2.5th/97.5th percentile). A feature was termed a robust age correlate if this 95% interval excluded zero.

Within each bootstrap iteration, raw p-values across all features were corrected for multiple comparisons using the Benjamini-Hochberg procedure; a feature’s selection probability is the proportion of the 1,000 iterations in which it was selected (p*<*0.05) after this correction.

A principal component analysis (PCA) was performed on the full set of robust features (z-scored) using the complete cohort. The full explained-variance spectrum across all components is reported. For the feature-contribution breakdown, only the leading components needed to reach 80% cumulative explained variance are shown. A feature’s squared loading on a component is reported as that feature’s percentage contribution to the component’s variance.

#### 5.5.2 Single-correlation analysis (fNIRS dataset)

The fNIRS-derived dataset’s small, unevenly distributed cohort (*N* = 13) does not support the stratified bootstrap described above. This dataset was treated as foundational, exploratory work rather than a basis for direct comparison with the other two datasets. For each feature, a single Pearson correlation with age was computed across the full cohort, reporting the raw correlation coefficient, the raw p-value, alongside the Benjamini-Hochberg-corrected p-value across all features tested.

#### 5.5.3 Cross-dataset comparison: age-correlate strength

Whether age correlates themselves are more strongly associated with age at one site than the other was assessed using the union of robust age correlates identified by the bootstrap procedure above at the temple (14 features) and wrist (3 features) sites, 15 unique features after accounting for 2 shared between sites. For each of the 15 union features, Cohen’s *d* was computed between the feature’s temple and wrist bootstrap distributions of *r*.

#### 5.5.4 Cross-dataset comparison: feature redundancy (“richness”)

To compare how redundant the feature sets were between the temple and wrist datasets, a matched stratified bootstrap was used: in each of 1,000 iterations, the same age-banding scheme was applied independently to both datasets. Within each iteration, the pairwise Pearson correlation matrix among all extracted features was computed separately for each dataset, and the mean absolute off-diagonal correlation was taken as a single richness value for that dataset in that iteration. This produced two paired bootstrap distributions of 1,000 richness values each. Cohen’s *d* quantified the effect size of level of feature “richness” between two sites.

## 6 Supplementary

## Notes

### Competing Interest Statement

All authors are full-time employees of Temple Wearables Inc, which funded this research. Temple Wearables Inc has commercial interests in wearable health technology and PPG based algorithms. The authors declare no other competing financial or non-financial interests

